# An enhanced-sampling MD-based protocol for molecular docking

**DOI:** 10.1101/434092

**Authors:** Andrea Basciu, Giuliano Malloci, Fabio Pietrucci, Alexandre M. J. J. Bonvin, Attilio V. Vargiu

## Abstract

Understanding molecular recognition of proteins by small molecules is key for drug design. Despite the number of experimental structures of ligand-protein complexes keeps growing, the number of available targets remains limited compared to the druggable genome, and structural diversity is generally low, which affects the chemical variance of putative lead compounds. From a computational perspective, molecular docking is widely used to mimic ligand-protein association *in silico*. Ensemble-docking approaches include flexibility through a set of different conformations of the protein obtained either experimentally or from computer simulations, e.g. molecular dynamics. However, structures prone to host (the correct) ligands are generally poorly sampled by standard molecular dynamics simulations of the apo protein. In order to address this limitation, we introduce a computational approach based on metadynamics simulations (EDES - <u>E</u>nsemble-<u>D</u>ocking with <u>E</u>nhanced-sampling of pocket <u>S</u>hape) to generate druggable conformations of proteins only exploiting their apo structures. This is achieved by defining a set of collective variables that effectively sample different shapes of the binding site, ultimately mimicking the steric effect due to ligands to generate holo-like binding site geometries. We assessed the method on two challenging proteins undergoing different extents of conformational changes upon ligand binding. In both cases our protocol generated a significant fraction of structures featuring a low RMSD from the experimental holo conformation. Moreover, ensemble docking calculations using those conformations yielded native-like poses among the top ranked ones for both targets. This proof of concept study paves the route towards an automated workflow to generate druggable conformations of proteins, which should become a precious tool for structure-based drug design.

## Introduction

Proteins are involved in virtually all cellular tasks and mediate physiological and pathological processes through the establishment of specific interactions with other biomolecules and small compounds. This feature is exploited in drug design whereby small molecules are developed to interfere with pathogenic pathways. Modern drug design relies on the detailed understanding of the molecular recognition process by which biological partners such as a protein and a drug interact and bind to each other^1–32^. From a structural perspective, the rapid increase in the number of experimentally-determined protein structures and the huge advances in computational resources have fueled the development of computer-aided strategies for drug design^4–7^. In particular, protein-ligand docking^8–10^ has become a well-established computational tool, often reducing the costs and improving the efficiency of high-throughput screenings. Docking algorithms provide a complementary alternative to experimental techniques such as X-ray crystallography, nuclear magnetic resonance, cryo-electron microscopy, and related methods for characterizing protein-ligand complexes^3^. However, as any computational or experimental technique, molecular docking also has its limitations and pitfalls, the treatment of partners’ flexibility being one of the most critical ones^1,3,11–17^. Indeed, molecular recognition is accompanied by various levels of structural changes occurring in both the ligand and the receptor. In proteins these changes go from relatively small side-chain rearrangements to local distortions involving loops and/or confined secondary structure variations, to even large-scale motions among different (sub)domains (e.g. hinge-bending or shearing motions)^2,18,19^. In particular, several classes of proteins including pharmaceutically relevant targets such as kinases^20^, transferases^21^, synthases^22^ and dehydrogenases^23^ undergo structural rearrangements leading to a compaction of the protein when bound to their substrates^24,25^.

In order to improve *in silico* structure-based drug design it is crucial to account for structural rearrangements (particularly those occurring at the binding site) when predicting drug binding and related thermodynamic and kinetic properties^3,8,9,26^. Unfortunately, most docking algorithms only consider limited receptor flexibility, often sampling a predetermined set of sidechains orientations and barely dealing with backbone rearrangements^1,9,12,17,27,28^. Such recipes often fail in predicting protein/ligand complexes in the presence of medium to large conformational changes of the receptor upon ligand binding. To cope with this issue, several methods have been developed over the last decades^3,5,28^, among which the so-called ensemble-docking has been shown to effectively enhance the performances of docking and virtual screening^4,5,11,12,29^. In a typical ensemble-docking calculation, different conformations of a protein target, either interacting with substrates other than those under study or free of any ligand, are used to improve the prediction of the correct structure of the complex of interest. The method is founded on the conformational selection / population shift theory of molecular recognition, stating that proteins are able to assume drug-bound (hereafter holo) like conformations even in the absence of interacting lig- and s^1,3,12,30^. The ligand thus recognizes its target by “selecting” the most complementary conformation from an ensemble of metastable states, causing a population shift toward holo-like states (structures).

The success of ensemble-docking is strongly dependent on the ability to include, in the pool of receptor structures, some conformation similar to the one found in the true complex^1,12,27,31–33^. In particular, it has been shown that the inclusion of experimental structures of proteins bound to ligands similar to the one of interest significantly increased the accuracy of the method^5,8,12,31,34^. However, with reference to the druggable genome^35^, the number of targets whose three-dimensional structure has been experimentally solved remains still limited^4^. Furthermore, the exploration of different conformations in experimental structures is generally limited and biased toward (often just a few) known ligand-receptor complexes, thus impacting on the chemical diversity of putative lead compounds in virtual screening campaigns. Computational methods including Monte Carlo and Molecular Dynamics (MD) offer a relatively cheap and complementary way to sample receptor conformations^4,5,11,12,17,36–40^. While the augmented conformational diversity sampled during MD simulations could in principle increase the percentage of false positives in virtual screening efforts (although this issue is also closely related to the limitations of current scoring functions^3,8,9,13,41,42^), the significance of including MD-derived structures for discovering new actives has been largely demonstrated^43,44^ e.g. by the discovery of new (sub)pockets not yet identified by experiments^45–49^. In fact, it has been proposed that MD-derived structures could capture key interacting spots on the surface of receptors that are less biased toward one specific chemotype^44^, potentially leading to the discovery of previously unknown activities and/or mechanisms of action (binding modes) of existing drugs^50^.

In the ensemble-docking framework, Lin et al.^51^ have introduced the concept of the Relaxed Complex Scheme (RCS) in which a series of independent docking runs are performed from receptor conformations of the unbound (hereafter apo) protein generated by MD simulations. A cluster analysis is generally performed to capture the structural diversity of the target (thus accounting for different functional (sub)states) while keeping the number of conformers to a computationally tractable number. Clearly, due to time scale restrictions, standard MD simulations are often unable to sample conformational states relevant to molecular recognition^38,52^. Therefore, several techniques have been proposed to enhance the sampling of rare conformations, including accelerated MD^53^, replica-exchange in temperature and energy spaces^54,55^, and metadynamics^56^, which generalizes methods such as conformational flooding^57^ and local elevation^58^. Several groups have demonstrated the power of these methods (sometimes coupled with the use of co-solvents) in improving the performance of docking and virtual screening^59–66^. Despite recent improvements however, no method has been developed yet that, without exploiting specific experimental information on the targets of interest, outperforms consistently standard ensemble docking when applied to targets undergoing different levels of conformational changes (from sidechain rearrangements to hinge-bending motions).

In order to address this issue, here we propose a new approach, EDES (<u>E</u>nsemble <u>D</u>ocking with <u>E</u>nhanced sampling of pocket <u>S</u>hape), which exploits relatively short metadynamics simulations of the apo protein of interest to generate a set of druggable (holo-like) conformations to be employed in ensemble-docking^56,67,68^. The key ingredients of our method are the use of a novel set of collective variables to sample in a controlled manner maximally different shapes of the binding site, and a multi-step clustering strategy allowing to retain a large fraction of holo-like structures within the pool of cluster representatives. Notably, EDES does not exploit specific information on the holo structure of the protein. We assess the method on two study cases representative of targets undergoing large and minor conformational rearrangements upon ligand binding (Figure 1 and Figure S1).

**Figure 1.**
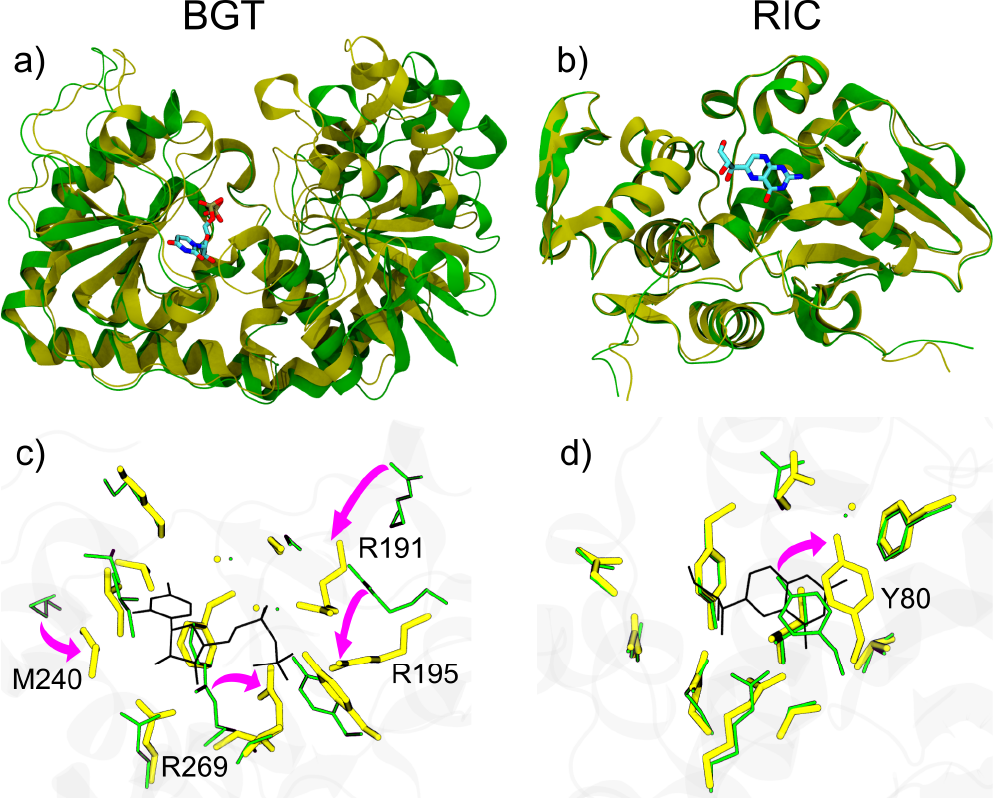
Comparison of the structural changes undergone by BGT (a-c) and RIC (b-d) upon binding of their ligands UDP and NEO, respectively. a-b) Rearrangements in the whole protein structure. The apo and holo proteins (PDB IDs 1JEJ, 1JG6 and 1RTC, 1BR5 for BGT, BGT-UDP and RIC, RIC-NEO systems respectively) are shown respectively in green and yellow ribbons, with the ligands in sticks colored by atom type; c-d) Zoom on local rearrangements at the binding site (BS). The conformations of residues lining the BS in the apo and holo forms of the proteins are shown in thin yellow and thick green sticks respectively, while the ligands are shown with thin black sticks and the protein in transparent grey ribbons. The most significant reorientations upon ligand binding are indicated by magenta arrows.

The first target is the T4 phage beta-glucosyltransferase (hereafter BGT)^69^, which displays a hinge-bending motion leading to a more closed form in its complex with uridine diphosphate (UDP) as compared to the ligand-free structure (Figure 1a,c). This protein was included in the set of 10 targets selected by Seeliger and de Groot^65^ to assess their workflow based on enhanced-sampling using tCONCOORD^70,71^ with the radius of gyration of the holo structure as a bias.

While close-to-native ligand binding poses were obtained for 8 out of 10 cases within the 100 top-ranked complex models, this was not the case for BGT, which makes this protein a well-suited test case for our method.

The second target is the recombinant ricin (hereafter RIC)72, representative of proteins undergoing minor but subtle conformational changes upon ligand (namely neopterin - NEO)73 binding (Figure 1 b,d). RIC belongs to the Astex Diverse Dataset74, recently used to validate the AutoDockFR docking software, which models receptor flexibility by explicitly specifying a set of side-chains for which rotatable bonds are active75. In cross-docking experiments using the apo conformations of the receptors, AutoDockFR out-performed AutoDock VINA76 in terms of number of correct poses and their ranking. However, none of the aforementioned software was able to find any solution within 2.5 Å (RMSD of the ligand) from the experimental structure of the complex.

In the following we demonstrate that for both these challenging targets EDES was able to generate native-like structures of the complexes. Moreover, using two wide-spread docking programs differing in search and scoring algorithms, namely HADDOCK and AutoDock4^77,78^, we identified native-like docking poses among the top ranked ones. While being a proof of concept, this work opens the way to the automatic generation of druggable conformations for a broad range of protein targets, and, as such, contributes to improving *in silico* structure-based drug design.

## Results and Discussion

In this section we first describe briefly the main workflow of EDES. Next, we demonstrate the effectiveness of the method in generating holo-like protein conformations. Finally, we ocompare our method with standard ensemble docking and with previous work.

### Method workflow

Our protocol workflow is sketched in Figure 2a. After identification of the putative binding site (hereafter BS - see Figure S1e for the list of residues defining those of BGT and RIC) on the target protein, we calculate the “inertia planes” at that site. These are defined as the planes orthogonal to the corresponding inertia axes and passing through the center of mass of the BS (Figure 2b). Then, we perform relatively short bias-exchange well-tempered metadynamics simulations^56,67,68^ of the apo protein (see Methods for details) using a set of four collective variables (hereafter CVs): three “(pseudo)contacts-across-inertia-planes” (hereafter CIPs) variables, each defined as the number of contacts between residues of the BS laying on opposite sides of the corresponding inertia plane (Figure 2c), and the gyration radius of the BS (RoG_BS_). We also use the latter CV to implement a “windows approach” (Figure 2d) aimed to sample more effectively and in a controlled manner different shapes of the BS (possibly mimicking conformational changes induced by ligand binding). Namely, in addition to the metadynamics bias applied on the 4 CVs, we apply soft restraints at values of the RoG_BS_ that are respectively 7.5% higher and lower compared to the value measured in the apo X-ray structure (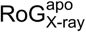, corresponding to the center of window 1). Next, from the trajectory corresponding to this first window, we randomly select a conformation of the protein whose RoG_BS_ is 5% lower than 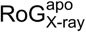 and perform another simulation (corresponding to window 2) with walls centered at ±7.5% 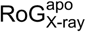 from this new center. We repeat this procedure up to four windows including the first one, which leads to an overall reduction of RoG_BS_ by 15% relatively to the center of the first window 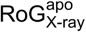 (see Figures 2, 3 and Table S1). Despite our choice is arbitrary, the performance of our protocol turns out to be not very sensitive to the number of windows chosen (thus, to the exact extent of collapse induced at the BS, *vide infra*). In particular, we obtained comparable results using either 3 or 4 windows. Moreover, the generality of our protocol was validated against RIC, which does not feature large conformational changes upon substrate binding.

**Figure 2.**
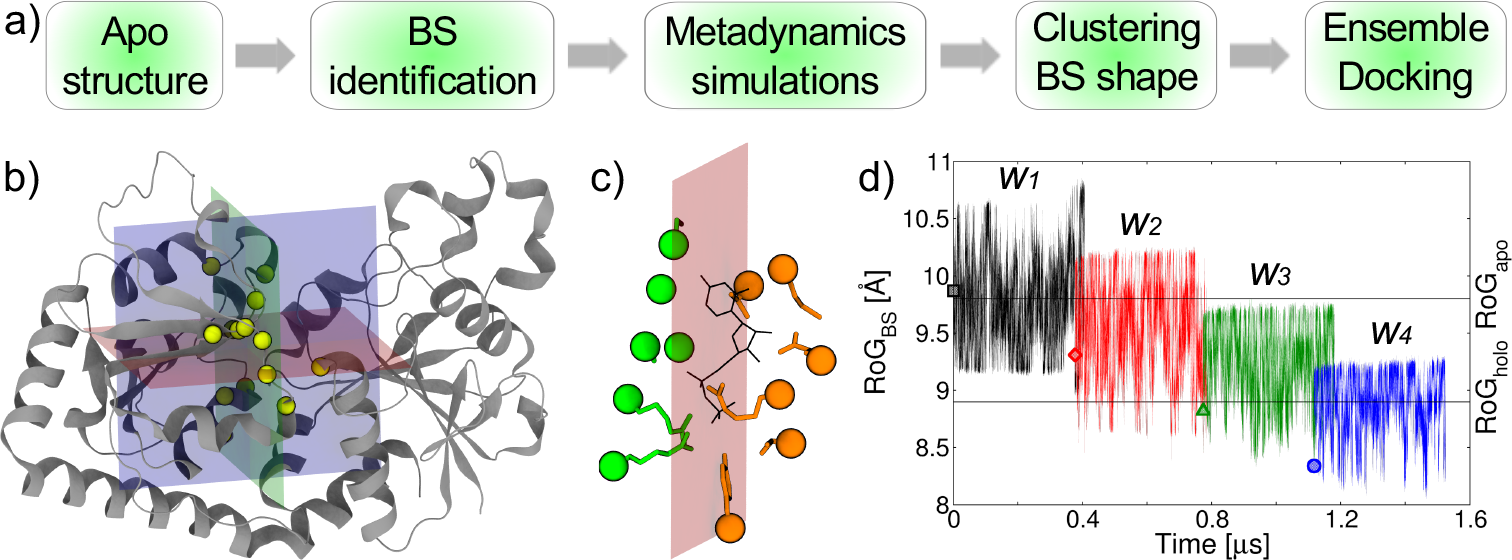
Overview of the EDES approach. a) Workflow of the EDES protocol; b) Representation of the “inertia planes” (transparent blue, red and green) calculated at the BS. Alpha carbons of residues lining this site are shown as yellow van der Waals spheres, while the protein is shown in white ribbons; c) Schematic view showing the definition of the two groups of atoms (orange and green sticks with alpha carbons as van der Waals spheres) considered for the calculation of the number of contacts across one inertia plane; the ligand is also shown in black sticks; d) Scheme of the “window approach” implemented to enhance in a controlled manner the sampling of conformations associated to different RoGBS values (the plot refers to simulations of the BGT system). For each of the (up to) four windows considered, the plot of RoGBS is shown with different colors. The values corresponding to the initial conformation for each window are shown by a square (w_1_), diamond (w_2_), upper triangle (w_3_) and circle (w_4_). The values of RoGBS calculated for the apo and holo experimental structures are also indicated by horizontal lines.

**Figure 3.**
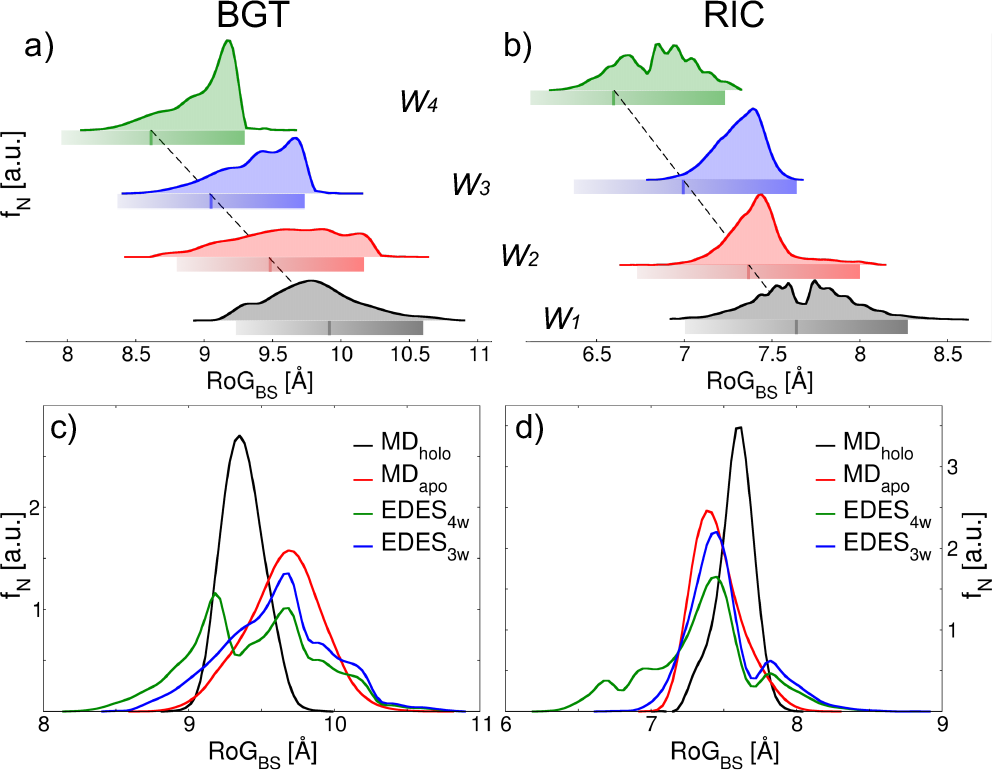
Distributions of RoGBS values. a-b) Distributions within each window of EDES for BGT (a) and RIC (b). The colored bars below each distribution indicate the position of the lower and upper walls set for RoGBS in that window, the color gradient indicating a higher force constant for the upper than the lower wall (the centers of the windows are indicated by a darker line within each bar and are connected by a virtual line - black dashed). c-d) Comparison of RoGBS normalized distributions (area under each curve equal to 1, bin size set to 0.1 Å) obtained from the different simulations performed in this work.

### Sampling of holo-like structures

Here we discuss the performance of our method in generating druggable holo-like structures of both targets. We start with BGT as it is a paradigm flexible protein undergoing hinge-bending motions upon binding of UDP^69^. The ligand induces large rearrangements at the BS particularly in the orientation of three arginines (R191, R195 and R269) neutralizing the negative charge of the diphosphate group (Figure 1a-c). We first compared the performance of standard MD simulations of the apo (MD_apo_) and holo (MD_holo_) systems to that of EDES (simulation details are reported in Methods and Table S1) using as metric the RMSD of the BS (RMSD_BS_) from the geometry assumed in the holo experimental structure. Figure 4 shows a very poor overlap between the MD_apo_ and MD_holo_ distributions, considering the data corresponding either to snapshots extracted from the MD simulations or the cluster structures. The EDES distributions are centered somewhat in between the ones obtained from unbiased MD simulations. In addition, most conformations feature an RMSD_BS_ lower than 2.8 Å (the value between the apo and holo experimental structures, see Figure S1e) from the experimental structure of the complex.

**Figure 4.**
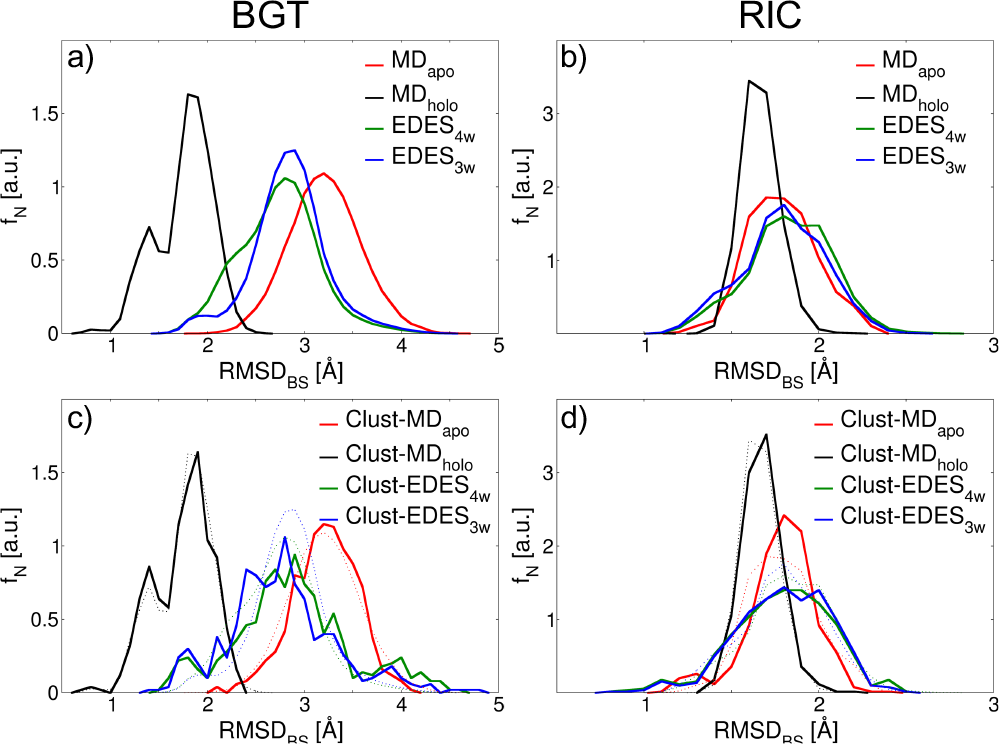
Normalized distributions (area under each curve equal to 1, bin size set to 0.1 Å) of RMSDBS with respect to the holo structure. a,c) Distributions for BGT obtained considering all snapshots extracted from MD simulations (a) and only from cluster representatives (c). Dotted tiny lines correspond to the distributions in a); b,d) same as in a,c) but for RIC.

Moreover, a prominent shoulder raises the percentage of conformations with RMSD_BS_ < 2 Å as compared to MD_apo_, a feature that persists also when inspecting the distributions obtained from the clusters sets (Figure 4 and Table 1). Interestingly, despite enhancing the sampling of the BS only, EDES is able to drag the whole protein structure towards conformations close to that found in the protein-ligand complex (Figure S3).

**Table 1.** Performance of various MD simulations in reproducing native-like conformations of the BS, measured as the percentage of conformations featuring a value of RSMD_BS_ lower than 1.5 or 2 Å. “Trajectory” and “Clusters” refer to snapshots extracted from the full trajectories and to the cluster analysis representatives, respectively. The lowest value of the RMSD (Å) with respect to the holo X-ray structures is reported in parentheses.

|  |  | RMSD<sub>BS</sub> < 1.5 Å [%] |  | RMSD<sub>BS</sub> < 2 Å [%] |  |
| --- | --- | --- | --- | --- | --- |
|  |  | Trajectory | Clusters | Trajectory | Clusters |
| BGT | MD<sub>apo</sub> | - | - | 0.06 (1.69) | - |
|  | MD<sub>holo</sub> | 23.9 (0.75) | 26.6 (0.75) | 85.3 (0.75) | 84.8 (0.75) |
|  | EDES<sub>4w</sub> | 0.02 (1.31) | 0.2 (1.31) | 5.0 (1.31) | 8.4 (1.31) |
|  | EDES<sub>3w</sub> | 0.02 (1.31) | 0.4 (1.31) | 3.8 (1.31) | 9.2 (1.31) |
| RIC | MD<sub>apo</sub> | 9.7 (1.00) | 9.6 (1.10) | 89.4 (1.00) | 93.2 (1.10) |
|  | MD<sub>holo</sub> | 13.0 (0.91) | 12.4 (0.95) | 99.6 (0.91) | 97.8 (0.95) |
|  | EDES<sub>4w</sub> | 13.0 (0.77) | 19.0 (0.81) | 81.4 (0.77) | 82.4 (0.81) |
|  | EDES<sub>3w</sub> | 16.5 (0.77) | 17.6 (0.81) | 85.5 (0.77) | 85.4 (0.81) |

Table 1 reports the percentage of structures with low RMSD_BS_ from the holo experimental structure. As expected, the percentage of such conformations is high for MD_holo_. Moreover, while a very low number of such conformations was sampled in MD_apo_, a consistent fraction was recorded by EDES using either 3 or 4 windows. In particular, by mimicking in part the hindrance of a ligand through the bias applied on the collective variables CIPs, our protocol was able to generate overall collapsed (particularly with respect to MD_apo_, see Figure S4) but “free-inside” conformations of the BS (Figure 5). Such behavior is particularly evident for R269, which is displaced towards one side of the BS (as it happens in the holo structure where this residue interacts with the negatively charged phosphate group of the ligand) only in EDES but not in MD_apo_. Moreover, our multi-step cluster analysis was able to effectively increase the percentage of structures featuring a native-like geometry of the BS with respect to the fraction sampled during MD simulations (Table 1). The enhanced sampling of holo-like conformations by EDES is evident also using the CIPs metric, as seen by the improved overlap between the MD_holo_ and EDES distributions as compared to MD_apo_ (Figure 6). In particular, only EDES is able to sample conformations featuring values of the CIPs variables virtually identical to those of the experimental holo structure (black sphere in Figure 6).

**Figure 5.**
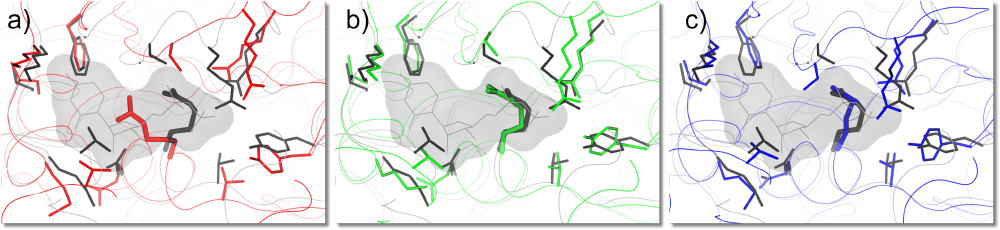
Conformations of BGT extracted from MD_apo_ (a), EDES_4w_ and EDES_3w_ (c) and associated to the lowest RMSD_BS_ with respect to the holo experimental structure. The molecular surface of UDP is shown in gray transparent color, with atom centers connected by solid lines. The proteins are shown as gray (experimental holo-form), red (MD_apo_), dark green (EDES_4w_) and blue (EDES_3w_) thin ribbons, with sidechains of residues lining the BS represented as sticks which are thicker for R269.

**Figure 6.**
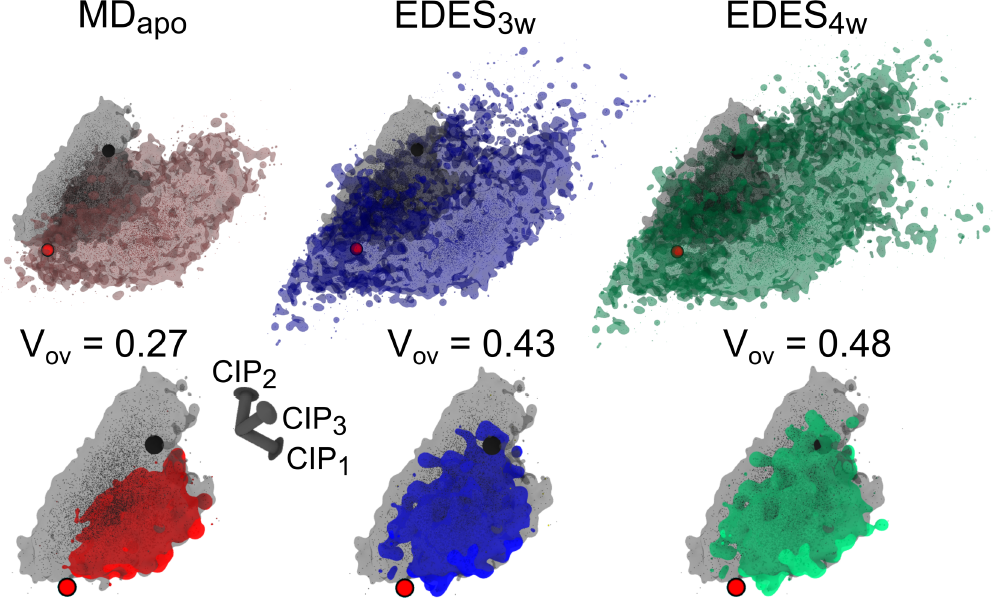
Sampling of the three-dimensional space defined by CIP_1-3_ during the MD simulations of BGT and BGT-UDP. Top row) Comparison of MD_holo_ (dark gray) distributions with MD_apo_ (red), EDES_3w_ (blue) and EDES_4w_ (green). Distributions are shown both as solid points and as transparent surfaces. The location of apo and holo structures are indicated by red and black spheres, respectively; bottom row) Envelopes of the overlapping portion between the distribution of MD_holo_ (shown in dark gray as reference) and those of MD_apo_ (red), EDES_3w_ (blue) and EDES_4w_ (green). The volumes of the overlapping distribution V_op_ are also reported (estimated with the Voss Volume Voxelator (3V) –http://3vee.molmovdb.org - using a probe radius of 3 Å).

Despite our method was primarily devised for flexible targets, in order to investigate its general applicability, we decided to validate it also on a protein undergoing minor conformational changes at the BS upon ligand binding in order to investigate how general could be its applicability. The recombinant ricin protein^72^ (RIC) is one of such targets, and was selected also because its subtle conformational changes upon binding of NEO (Figure 1b,d) were hardly handled by algorithms exploiting flexibility of the BS only in terms of activation of sidechain torsionals^75^. In particular, RIC resulted a very difficult target either for rigid or flexible docking calculations performed on apo X-ray structures with AutoDock VINA and the recently introduced AutoDockFR software (see Table 1 in ^75^). Ensemble-docking approaches (despite being computationally demanding compared to flexible docking on single structures) were on the contrary able to reproduce the correct structure assumed by the BS in the holo structure (Table 1).

In this case, the performances of standard and enhanced-sampling MD simulations are overall similar (although EDES was able to find BS conformations closer to the holo experimental structure than those obtained from MD_apo/holo_, see Table 1 and Figure 4).

As expected, in this case there is also no clear difference between EDES and standard MD in reproducing holo-like conformations of the protein (Figure S3), both approaches being able to sample a relatively large fraction of such structures. On the basis of these results, we are confident that our approach, although originally devised for proteins undergoing extended conformational changes, is effective in generating holo-like structures also of targets undergoing minor conformational changes upon binding. This is particularly important since in a real case one might not know the extent of the conformational change in advance.

### Docking performance

In this subsection we describe the performance of each set of structural clusters in ensemble-docking calculations. Regarding BGT, both AutoDock4 and HADDOCK displayed an improved sampling performance (defined as the percentage of docking poses displaying a value of RMSD_lig_ lower than 2 Å from the ligand conformation in the holo experimental structure) when coupled to EDES rather than MD_apo_ (Tables 2, 3). Namely, our approach was able to generate a consistent fraction (up to 2% and 14% with AutoDock4 and HADDOCK, respectively) of native-like ligand poses, performing much better than when starting from the clusters derived from (the much longer) MD_apo_ (no - AutoDock4 –, or 2% - HADDOCK - of native-like poses). Importantly, both programs were able to rank a native-like pose among the top three according to their respective clustering, scoring, and pose selection schemes when coupled with EDES, independently on the number of windows used to generate conformational clusters (see Tables 2, 3 and Figure 7).

**Table 2.** Performance of AutoDock4 in reproducing the experimental structures of the BGT-UDP and RIC-NEO complexes in ensemble-docking calculations. Results refer to clusters of ligand poses (500 for each ensemble of clusters of receptor structures, corresponding to the top pose from each independent docking run for that ensemble) generated using a distance matrix metrics (dRMSD) with a cutoff of 1.5 Å. The sampling performance is calculated as the percentage of poses within 2 Å from the native structure out of the 500 top poses considered for each ensemble of receptor structures. The fourth row reports the ranking of the first native-like pose obtained using the highest score with-in each cluster for ranking. In parentheses, the rank of the same cluster is reported when the average score over the top three poses is used instead. The fifth row reports the population of the corresponding cluster in the same column. The last row reports the average heavy-atoms RMSD of the ligand calculated for the top cluster, with standard deviation in parentheses.

|  | BGT |  |  |  | RIC |  |  |  |
| --- | --- | --- | --- | --- | --- | --- | --- | --- |
|  | MD<sub>apo</sub> | MD<sub>holo</sub> | EDES<sub>3w</sub> | EDES<sub>4w</sub> | MD<sub>apo</sub> | MD<sub>holo</sub> | EDES<sub>3w</sub> | EDES<sub>4w</sub> |
| **Sampling performance [%]** | - | 84.6 | 1.8 | 2.0 | 3.8 | 10.8 | 2.6 | 3.4 |
| **Cluster rank** | - | 1 (1) | 1 (1) | 2 (2) | 1 (1) | 1 (1) | 4 (3) | 1 (1) |
| **Cluster population** | - | 48 | 4 | 4 | 15 | 20 | 13 | 13 |
| **RMSD<sub>lig</sub> [Å]** | - | 1.2 (0.7) | 0.6 (0.9) | 1.5 (1.2) | 1.0 (0.9) | 0.6 (0.6) | 0.7 (0.7) | 0.9 (0.8) |

**Table 3.** Performance of HADDOCK in reproducing the experimental structures of the BGT-UDP and RIC-NEO complexes in ensemble-docking calculations with different sets of protein conformational clusters. Results correspond to the statistic of the top pose in clusters obtained using a ligand interface RMSD metrics with a 1 Å cutoff and minimum number of 4 poses per cluster. The cluster rankings are based on the average score of the top 4 poses (the ranking based on the score of the top pose is reported between brackets). In the last two rows the first values reported refer to the average statistics of the top 4 poses of a cluster in the semi-flexible refinement (it1 step) of HADDOCK, while values in parentheses refer to the statistics/rank of the best (smallest RMSD to reference) pose in the top 4. The sampling performance is calculated as the percentage of poses within 2 Å from the experimental structure out of the 400 generated models (one docking run was performed from the ensemble of 500 MD conformation). F_nat_ indicates the fraction of native contacts recovered within a shell of 5 Å from the ligand in the experimental structures. See Table 2 for further details.

|  | BGT |  |  |  | RIC |  |  |  |
| --- | --- | --- | --- | --- | --- | --- | --- | --- |
|  | MD<sub>apo</sub> | MD<sub>holo</sub> | EDES<sub>3w</sub> | EDES<sub>4w</sub> | MD<sub>apo</sub> | MD<sub>holo</sub> | EDES<sub>3w</sub> | EDES<sub>4w</sub> |
| **Sampl. Perf. [%]** | 1.8 | 17.5 | 14.0 | 8.0 | 23.0 | 45.5 | 15.8 | 20.5 |
| **Pose rank** | 19 (126)<sup>a</sup> | 1 (1) | 2 (5) | 1 (6) | 1 (1) | 1 (2) | 1 (1) | 1 (1) |
| **Clus. Pop.** | - | 9 | 33 | 5 | 80 | 152 | 53 | 69 |
| **F<sub>nat</sub>** | 0.75 (0.62)<sup>a</sup> | 0.81±0.07 (0.83/1) | 0.80±0.02 (0.79/3) | 0.72±0.04 (0.71/3) | 0.93 (0.93/1) | 0.90±0.03 (0.87/4) | 0.90±0.06 (1.0/3) | 0.83±0.06 (0.73/3) |
| **RMSD<sub>lig</sub> [Å]** | 1.8 (1.4)<sup>a</sup> | 1.6±0.6 (0.86/1) | 1.1±0.3 (0.7/3) | 1.9±0.4 (1.5/3) | 1.3±0.3 (0.95/1) | 1.2±0.2 (0.8/4) | 0.9±0.3 (0.4/3) | 0.9±0.2 (0.5/3) |

a. Since no acceptable clusters were obtained in this case, the reported statistics correspond to single pose statistics for the first acceptable ( $< 2 \text{ \AA}$ ) and best (between brackets) poses.

**Figure 7.**
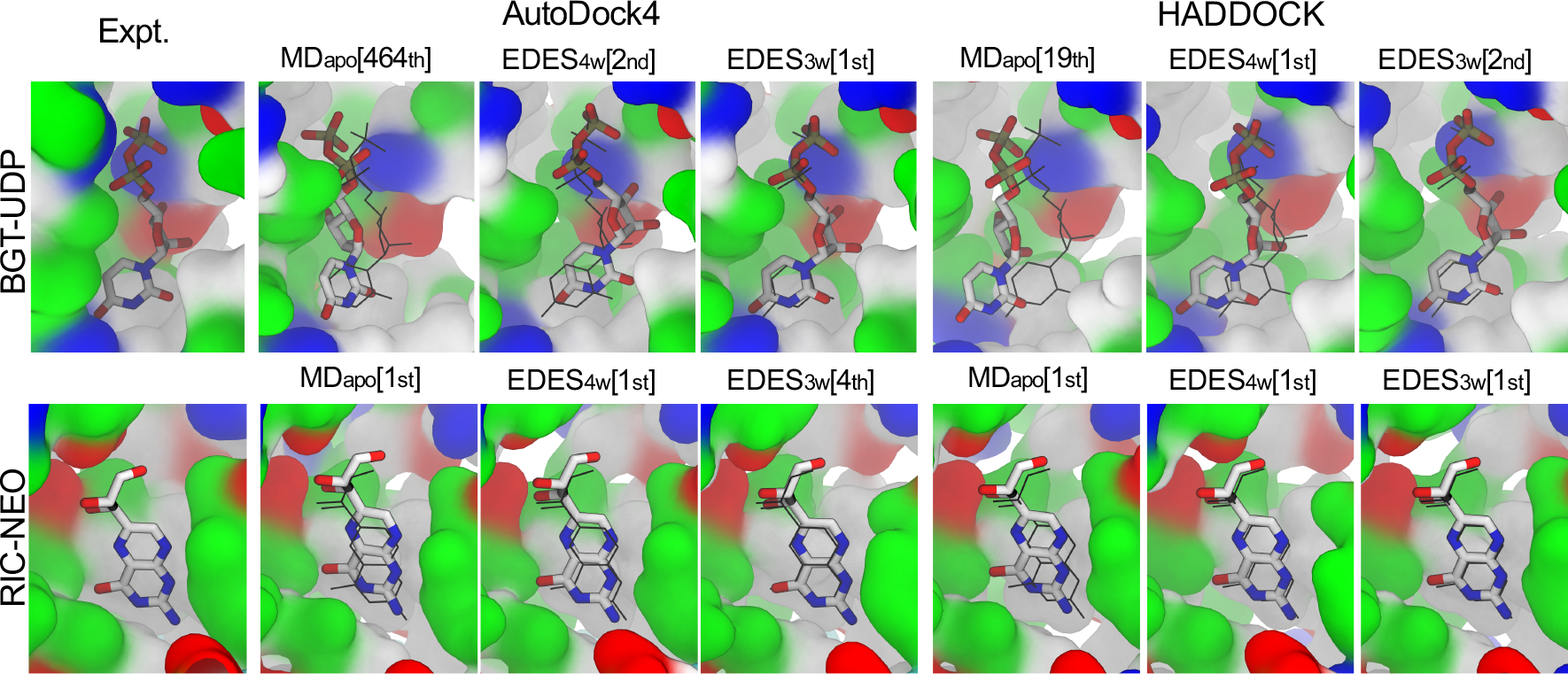
Docking performance of various structural ensembles in reproducing the experimental poses of UDP (top row) and RIC (bottom row). A closed view of the binding within the experimental BS is shown in the first column, while the top-score pose within the first cluster with RMSD_lig_ < 2 Å, or the pose with the lowest RMSD_lig_ value when no native-like pose was found, are reported in the next columns, with corresponding ranks in square brackets. The docking was performed both using AutoDock4 and HADDOCK for comparison. The molecular surface of the backbone and of the *C_α_* atoms of the protein are colored by residue type as in Figure 1, and the ligand is shown as sticks colored by atom type. In columns 2 to 7 the experimental pose is shown in black thin sticks for easy comparison.

Regarding RIC, as expected from the results discussed above, also the sampling performance of both AutoDock4 and HADDOCK increased significantly with respect to BGT using the set of cluster structures obtained from MD_apo_ or EDES (Tables 2, 3). Most importantly, also for RIC, EDES was able to rank native-like ligand poses among the top ones (Tables 2, 3 and Figure 7). This finding, therefore, extends the initial scope of our methodology to a different class of targets undergoing minor structural rearrangement upon ligand binding.

### Comparison with previous work

It is instructive to compare the performance of our method to previous computational work on the same target proteins. In ^65^ the tCONCOORD^70,71^ method was used to enhance the sampling of holo-receptor conformations of a set of 10 proteins including BGT, using the gyration radii of the holo proteins as bias. In 9 out of 10 cases the models generated by tCONCOORD featured an RMSD of the BS (defined there by the list of residues within 6 Å from the ligand in the experimental structure - hereafter BS6) smaller than 2 Å. In particular, the best model for BGT had an RMSD_BS6_ of 1.78 Å^65^, significantly higher than the lowest values obtained with our protocol, namely 1.51 Å and 1.40 Å for the clusters and the MD-derived distributions in both EDES_4w_ and EDES_3w_. In 65 ensemble-docking calculations with AutoDock VINA76 were performed on 5000 protein structures generated by tCONCOORD, followed by a series of post-docking optimizations, filtering of models against the experimental gyration radius, further docking calculations and rescoring with RosettaLigand^79^.

As a result, in 8 out of 10 cases native-like ligand poses (defined there as those for which RMSD_lig_ < 3 Å with respect to the experimental structure) were generated among the top 100 ones, demonstrating the general applicability of the method for blind predictions of protein-ligand complexes involving large conformational rearrangements. However, no native-like pose was found within the top 100 ones for BGT, and just one pose featured an RMSD_lig_ < 2 Å.

In order to understand more deeply the reasons behind the good performance of our method, we calculated the correlation between RMSD_lig_ and the RMSDs of various residues selections: BS6, BS, the arginine triad (R191, R195 and R269), and R269, which in the MD_apo_ simulation often occupies the center of the BS (Figure 5). The results indicated that, while the overall correlation between RMSD_UDP_ and the RMSD of BS (and even more of BS6) is not necessarily high, reproducing the correct orientation of the arginine triad and in particular of R269 is crucial to obtain native-like poses of UDP (Figure S5). Thus, by enhancing the fluctuations in the number of contacts among two relatively small groups of atoms across three orthogonal planes (see e.g. Figure 6 and Figure S4), our method effectively induces the “accessible space” within the BS to assume different volumes and shapes, increasing in this way the probability of sampling native-like conformations. This should be particularly effective when dealing with long sidechain charged residues lining the BS (such as R269) within an aqueous environment that favors an extended sidechain conformation due to enhanced hydration. This, in combination with the controlled bias applied on RoG, allows to obtain “open-in-the-middle” conformations of an otherwise relatively closed BS, as it happens in the experimental structure of the complex. Regarding RIC, as stated above in ^75^ neither AutoDock VINA nor the therein introduced AutoDockFR docking software were able to find native-like poses of NEO. In particular, RIC showed to be a very difficult target for both rigid and flexible docking calculations with 7 rotatable bonds for the ligand and 7 flexible sidechains in the protein (see Table 1 in ^75^). In contrast, our protocol performed very well in reproducing holo-like structures. Moreover, the performance of EDES was similar to that of ensemble docking using structures from MD_apo_. This is not trivial, as recently discussed in ^62^ where the advantage of using an enhanced-sampling protocol (namely accelerated MD^53^) vs. conventional MD simulations was reported to depend on the target, in particular on the extent of conformational changes at the BS and on the binding specificity. Our findings for RIC are thus very encouraging considering the difference of only 0.1 Å between the RoG_BS_ of the apo and holo experimental structures (see Figure 1) and the relatively large fluctuations induced at the BS by our protocol vs. those induced by standard MD simulations (Figure 3).

### Concluding Remarks

We have presented a proof of concept study of a novel protocol for ensemble-docking. Our approach was able to generate a relevant fraction of holo-like conformations of the proteins and rank the native-like ligand poses among the top ones. Its robustness and general applicability were tested using two different docking programs against two challenging protein targets undergoing different extents of conformational changes upon ligand binding. The two key points of our method are: *i)* the use of soft adaptive biases on a carefully designed new set of CVs, enabling the generation of maximally diverse conformations of the BS, including a relevant fraction of holo-like ones. A crucial feature of this set is related to its ability to produce “coconut-like” conformations of the BS, that is geometries that are ligand-accessible despite being relatively shrunk. As such, we infer that our protocol will generate druggable conformations of proteins featuring a partial collapse of the BS upon binding; *ii)* a multi-step cluster analysis performed on the CVs able to generate a tractable number of conformations while maintaining or even increasing (with respect to distributions extracted from MD simulations) the fraction of holo-like structures.

In perspective, a straightforward way to further improve the sampling of different (and druggable) conformations of the BS could be coupling our algorithm to co-solvent simulations^66,80^, as done e.g. in ^63^. Furthermore, our method could be combined with others enhancing the sampling of orthogonal degrees of freedom, such as global protein motions^60,81^, rotations around torsional angles^61,62^, secondary structure changes^82,83^, rescaled protein-ligand interactions^54,63^, just to cite a few options. In addition, experimental information from many sources could be easily encoded in new CVs and/or restraints. We plan to extend the method so as to sample also expanded conformations of the BS (in order to deal with non-specific protein targets such as the acetylcholine binding protein displaying opening or closing of the site upon binding of different ligands^62^). Note however that already in the current implementation our protocol was able to generate a fraction of such structures for the test-cases considered in this work (Figures 3, 6 and Figure S4). As a long-term goal, we aim to create a database of proteins structures that should help in reducing the cost associated to the generation of the structures for ensemble-docking runs, allowing for a single target virtual screening of thousands of compounds in a reasonable amount of time. In this perspective, the ensemble of targets could be also used to repositioning existing drugs for new therapeutic uses as recently shown^50^.

## Materials and Methods

### Standard MD simulations

Standard all-atom MD simulations were carried out using the *pmemd* module of the AMBER16^84^ molecular modeling software. Topology files were created for each system using the LEaP module of AmberTools17 and starting from the experimental structures available in the PDB databank (PDB IDs 1JEJ, 1RTC, 1JG6 and 1BR5 for BGT, BGT-UDP, RIC and RIC-NEO systems respectively)^69,72,73^. The ff14SB^85^ and GAFF^86^ force fields were used for the proteins and the ligands, respectively. Missing parameters for the latter were generated using the antechamber module of AmberTools17. In particular, atomic restrained electrostatic potential charges were derived after a structural optimization performed with Gaussian09^87^. Each structure was solvated with explicit TIP3P water model, and its net charge was neutralized with the required number of randomly placed K^+^/Cl^-^ ions. The total number of atoms was ~86.000 for BGT and ~54.000 for RIC. Periodic boundary conditions were employed with long-range electrostatic as evaluated through the particle-mesh Ewald algorithm using a real-space cutoff of 12 Å and a grid spacing of 1 Å per grid point in each dimension. The van der Waals interactions were treated by a Lennard–Jones potential, using a smooth cutoff (switching radius 10 Å, cutoff radius 12 Å). The initial distance between the protein and the edge of the box was set to be at least 16 Å in each direction. Multi-step energy minimization with a combination of steepest descent and conjugate gradient methods was carried out to relax internal constrains of the systems by gradually releasing positional restraints. Following this, the systems were heated from 0 to 310 K in 10 ns of constant pressure heating (NPT) using the Langevin thermostat (collision frequency of 1 ps^−1^) and the Berendsen barostat. After equilibration, four production runs of 2.5 μs each were performed for the apo systems, while a single 1 μs-long simulation was performed for each complex. A time step of 2 fs was used for pre-production runs, while equilibrium MD simulations were carried out with a time step of 4 fs in the NPT ensemble (using a Monte Carlo barostat) conditions after hydrogen mass repartitioning^88^. Coordinates from production trajectories were saved every 100 ps and 10 ps for MD_apo_ and MD_holo_ respectively.

### Metadynamics simulations

Bias-exchange well-tempered metadynamics simulations^56,67,68^ were performed on the two apo proteins using the GROMACS 2016.5 package^89^ and the PLUMED 2.3.5 plugin^90^. The starting structure for each simulation was the last conformation saved from the equilibration step from MD_apo_. AMBER parameters were ported to GROMACS using the acpype parser^91^. To enhance the sampling of different BS shapes, we employed the following four CVs defined by including all heavy atoms of the residues lining the BS itself (here defined by the residues lining within 3 Å from the ligand in the experimental structure of the complex, see Figure S1 for the full lists for BGT and RIC): the radius of gyration of the BS (RoG_BS_), calculated using the “*gyration*” built-in function of PLUMED; the number of (pseudo)contacts across the “inertia planes” (CIP_1,2,3_) of the BS, defined as the planes orthogonal to each principal inertia axes and passing through the center of mass of the BS. These CVs were calculated by an *in-house* tcl script based on VMD *orient* function. Namely, residues lining the BS were split into two lists A and B according to the position of the geometrical center of their backbone on each of the two sides of the inertia plane, and the overall number of pseudo-contacts N_c_ between the two groups was calculated through the “*coordination*” keyword of PLUMED, which implements a switching function such as the following:

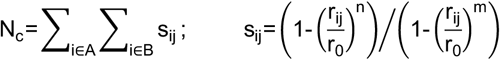

with r_0_ = 8 Å, n = 6, m = 12. Each replica was simulated for 100 ns (note that our aim is primarily to enhance sampling of different shapes of the BS and not to obtain converged free energy profiles), so that each window cumulated 400 ns of simulation time. The height *w* was set to 0.6 kcal/mol for both systems, while the widths *s_i_* of the Gaussian hills were set according to established prescriptions^92^ to 0.15 Å and 0.05 Å (RoG_BS_), 5.4 and 4.8 (CIP_1_), 5.1 and 3.2 (CIP_2_), 5.3 and 3.1 (CIP_3_) for BGT and RIC respectively. Hills were added every 2 ps, while the bias-exchange frequency was set to 20 ps. The bias factor for well-tempered metadynamics was set to 10. The “windows approach” briefly described in Results and Discussion was implemented using RoG_BS_ as control parameter. Namely, we applied restraints (force constants set to 50 and 10 kcal mol^−1^ Å^−2^ for upper and lower walls, respectively, as we seek for compression rather than enlargement of the BS) at values of the RoG_BS_ that are respectively 7.5% higher and lower compared to the value measured in the apo X-ray structure 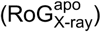. Then, from the trajectory corresponding to this first window, we select a random conformation of the protein whose RoG_BS_ is 5% lower than 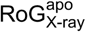 and perform another simulation with walls centered at ±7.5% 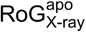 from this new center, repeating this procedure so as to simulate a total of four windows (see Figure 3 and Table S1). Note that the walls were set to allow partial overlap between adjacent windows, which indeed occurred in all cases (Figure 3).

### Cluster analysis of MD trajectories

The cluster analysis was performed on the four CVs defined above using R scripts developed *in house*. We implemented a multi-step strategy aimed at increasing in an unbiased manner the percentage of conformations similar to the native structure of the holo protein. Namely, the distribution of RoG_BS_ values sampled during the MD simulation was binned into 10 equally-wide slices, and a hierarchical agglomerative clustering (using the built-in function “hclust” and setting to “Euclidean” the method to compute the distance matrix) was performed on the four CVs within each slice, setting the number of generated clusters to *x_i_* = *N_i_/N_tot_* · *N_c_*, where *N_i_*, *N_tot_* and *N_c_ = 500* are the number of structures within the *i^th^* slice, the total number of structures, and the total number of clusters respectively. The resulting *N_c_* clusters were used as starting point to perform a second cluster analysis with the K-means method and requiring a total of *N_c_* clusters (maximum number of iterations set to 10000). Despite not making any use of specific knowledge of the structure of the complexes, our informed strategy was able to generate a larger fraction of cluster structures displaying an RMSD_BS_ < 2 Å than that obtained from the standard application of K-means using randomly selected conformations as starting points (Figure S2). In fact, this latter initialization strategy is considered one of the most unreliable ones based on a comparison of several alternative algorithms on a range of diverse data sets^93^.

### Molecular docking

Molecular docking calculations were performed with AutoDock4^78^ and the HADDOCK web server version2.2^77,94^ This choice allowed to validate our methodology against two programs differing in search algorithms, scoring functions, and pose selection schemes. Both programs were first validated for redocking against experimental structures (Table S2). Next, they were used to perform guided docking (see Figure S1 for the definition of the BS) with their default settings, apart from the following changes. In AutoDock4, the grid density (*spacing* parameter changed from 0.375 Å to 0.25 Å), and number of energy evaluations (*ga_num_evals* increased by a factor of 10 from the default value) were both increased, with the purpose to avoid repeating each calculation several times to obtain converged results. For each set of structures, 500 rigid docking independent calculations were performed using an adaptive grid enclosing all the residues belonging to the BS. Next, the top poses (in total 500, one for each docking run) were clustered using the *cpptraj* module of AmberTools17 with a hierarchical agglomerative algorithm and a cutoff of 1.5 Å for the RMSD distance matrix. For HADDOCK, a single docking run was performed per case, starting from the various ensembles of 500 conformations, with increased sampling (10000/400/400 models for rigid body docking, semi-flexible refinement and final refinement in explicit solvent). The weight of the intermolecular van der Waals energy for the initial rigid-body docking stage was increased to 1.0 (from the default 0.01), RMSD-based clustering was selected with a cutoff of 1 Å and the docking was guided by ambiguous distance restraints defined for the residues of the BS (Figure S1e) and the ligand as described in ^95^. In the rigid-body stage the protein BS residues were defined as “active”, effectively drawing the ligand into the BS without restraining its orientation. For the subsequent stage the restraints were such that only the ligand was active, allowing it to explore better the BS while maintaining at least one contact with its residues.

### Figures and graphs

Figures were generated with Maestro^96^, VMD 1.9.3^97^ and InkScape 0.91. Graphs were created with xmgrace 5.1.25.

## Author Contributions

Andrea Basciu and Attilio V. Vargiu designed research with contributions of all authors. Andrea Basciu performed molecular dynamics simulations. Alexandre M. J. J. Bonvin performed docking calculations with HADDOCK. Attilio V. Vargiu performed docking calculations with AutoDock4. All the authors contributed analysis tools and analyzed the data. The manuscript was written through contributions of all authors. All authors have given approval to the final version of the manuscript.

## Acknowledgment

We thank Stefano Forli (Scripps Research Institute, La Jolla), Arianna Fornili (Queen Mary University, London), Silvia Marchesan (University of Trieste) and Alessandro Pandini (Brunel University, London) and for the critical reading of the manuscript. Andrea Basciu, Giuliano Malloci and Attilio Vargiu are grateful to Andrea Bosin and Giovanni Serra (University of Cagliari) for helping with software installation and optimization, and to Paolo Ruggerone (University of Cagliari) for his continuous support. This work has been done as part of the BioExcel CoE (www.bioexcel.eu), a project funded by the European Union contract H2020EINFRA-2015-1-675728. The research leading to the results discussed here was partly conducted as part of the Translocation Consortium (http://www.translocation.eu) and has received support from the Innovative Medicines Initiative Joint Undertaking under Grant Agreement no. 115525, resources that are composed of financial contribution from the European Union’s Seventh Framework Programme (FP7/2007-2013) and EFPIA companies in kind contribution. The authors declare no competing financial interest.

## References

(1) Antunes, D. A.; Devaurs, D.; Kavraki, L. E. Understanding the Challenges of Protein Flexibility in Drug Design. Expert Opin. Drug Discov. 2015, 10 (12), 1301–1313.

(2) Babine, R. E.; Bender, S. L. Molecular Recognition of Protein−Ligand Complexes: Applications to Drug Design. Chem. Rev. 1997, 97 (5), 1359–1472.

(3) Du, X.; Li, Y.; Xia, Y.-L.; Ai, S.-M.; Liang, J.; Sang, P.; Ji, X.-L.; Liu, S.-Q. Insights into Protein-Ligand Interactions: Mechanisms, Models, and Methods. Int. J. Mol. Sci. 2016, 17 (2), 144.

(4) De Vivo, M.; Masetti, M.; Bottegoni, G.; Cavalli, A. Role of Molecular Dynamics and Related Methods in Drug Discovery. J. Med. Chem. 2016, 59 (9), 4035–4061.

(5) Ganesan, A.; Coote, M. L.; Barakat, K. Molecular Dynamics-Driven Drug Discovery: Leaping Forward with Confidence. Drug Discov. Today 2017, 22 (2), 249–269.

(6) Pagadala, N. S.; Syed, K.; Tuszynski, J. Software for Molecular Docking: A Review. Biophys. Rev. 2017, 9 (2), 91–102.

(7) Śledź, P.; Caflisch, A. Protein Structure-Based Drug Design: From Docking to Molecular Dynamics. Curr. Opin. Struct. Biol. 2018, 48, 93–102.

(8) Ferreira, L.; dos Santos, R.; Oliva, G.; Andricopulo, A. Molecular Docking and Structure-Based Drug Design Strategies. Molecules 2015, 20 (7), 13384.

(9) Forli, S. Charting a Path to Success in Virtual Screening. Molecules 2015, 20 (10), 18732.

(10) Irwin, J. J.; Shoichet, B. K. Docking Screens for Novel Ligands Conferring New Biology. J. Med. Chem. 2016, 59 (9), 4103–4120.

(11) Amaro, R. E.; Baudry, J.; Chodera, J.; Demir, Ö.; McCammon, J. A.; Miao, Y.; Smith, J. C. Ensemble Docking in Drug Discovery. Biophys. J. 2018, 114 (10), 2271–2278.

(12) Buonfiglio, R.; Recanatini, M.; Masetti, M. Protein Flexibility in Drug Discovery: From Theory to Computation. ChemMedChem 2015, 10 (7), 1141–1148.

(13) Chen, Y.-C. Beware of Docking! Trends Pharmacol. Sci. 2015, 36 (2), 78–95.

(14) Grinter, S.; Zou, X. Challenges, Applications, and Recent Advances of Protein-Ligand Docking in Structure-Based Drug Design. Molecules 2014, 19 (7), 10150.

(15) Kastritis, P. L.; Bonvin, A. M. J. J. On the Binding Affinity of Macromolecular Interactions: Daring to Ask Why Proteins Interact. J. R. Soc. Interface 2013, 10 (79).

(16) Moitessier, N.; Englebienne, P.; Lee, D.; Lawandi, J.; Corbeil, C. R. Towards the Development of Universal, Fast and Highly Accurate Docking/Scoring Methods: A Long Way to Go. Br. J. Pharmacol. 2008, 153 (S1), S7–S26.

(17) Zacharias, M. Accounting for Conformational Changes during Protein-Protein Docking. Curr. Opin. Struct. Biol. 2010, 20 (2), 180–186.

(18) Lexa, K. W.; Carlson, H. A. Protein Flexibility in Docking and Surface Mapping. Q. Rev. Biophys. 2012, 45 (3), 301–343.

(19) Wei, G.; Xi, W.; Nussinov, R.; Ma, B. Protein Ensembles: How Does Nature Harness Thermodynamic Fluctuations for Life? The Diverse Functional Roles of Conformational Ensembles in the Cell. Chem. Rev. 2016, 116 (11), 6516–6551.

(20) Wu, P.; Nielsen, T. E.; Clausen, M. H. FDA-Approved Small-Molecule Kinase Inhibitors. Trends Pharmacol. Sci. 2015, 36 (7), 422–439.

(21) Arrowsmith, C. H.; Bountra, C.; Fish, P. V.; Lee, K.; Schapira, M. Epigenetic Protein Families: A New Frontier for Drug Discovery. Nat. Rev. Drug Discov. 2012, 11, 384.

(22) Richard Flavin; Stephane Peluso; Paul L Nguyen; Massimo Loda. Fatty Acid Synthase as a Potential Therapeutic Target in Cancer. Future Oncol. 2010, 6 (4), 551–562.

(23) Che-Hong Chen; Julio Cesar Batista Ferreira; Eric R. Gross; Daria Mochly-Rosen. Targeting Aldehyde Dehydrogenase 2: New Therapeutic Opportunities. Physiol. Rev. 2014, 94 (1), 1–34.

(24) Amaral, M.; Kokh, D. B.; Bomke, J.; Wegener, A.; Buchstaller, H. P.; Eggenweiler, H. M.; Matias, P.; Sirrenberg, C.; Wade, R. C.; Frech, M. Protein Conformational Flexibility Modulates Kinetics and Thermodynamics of Drug Binding. Nat. Commun. 2017, 8 (1), 2276.

(25) Hayward, S. Identification of Specific Interactions That Drive Ligand-Induced Closure in Five Enzymes with Classic Domain Movements. J. Mol. Biol. 2004, 339 (4), 1001–1021.

(26) Fischer, M.; Coleman, R. G.; Fraser, J. S.; Shoichet, B. K. Incorporation of Protein Flexibility and Conformational Energy Penalties in Docking Screens to Improve Ligand Discovery. Nat. Chem. 2014, 6, 575.

(27) Riccardo Baron; J. Andrew McCammon. Molecular Recognition and Ligand Association. Annu. Rev. Phys. Chem. 2013, 64 (1), 151–175.

(28) Yuriev, E.; Holien, J.; Ramsland, P. A. Improvements, Trends, and New Ideas in Molecular Docking: 2012-2013 in Review. J. Mol. Recognit. 2015, 28 (10), 581–604.

(29) Huang, S.-Y.; Zou, X. Ensemble Docking of Multiple Protein Structures: Considering Protein Structural Variations in Molecular Docking. Proteins 2007, 66 (2), 399–421.

(30) Boehr, D. D.; Nussinov, R.; Wright, P. E. The Role of Dynamic Conformational Ensembles in Biomolecular Recognition. Nat. Chem. Biol. 2009, 5, 789.

(31) Korb, O.; Olsson, T. S. G.; Bowden, S. J.; Hall, R. J.; Verdonk, M. L.; Liebeschuetz, J. W.; Cole, J. C. Potential and Limitations of Ensemble Docking. J. Chem. Inf. Model. 2012, 52 (5), 1262–1274.

(32) Moroni, E.; Paladino, A.; Colombo, G. The Dynamics of Drug Discovery. Curr. Top. Med. Chem. 2015, 15 (20), 2043–2055.

(33) Wong, C. F. Flexible Receptor Docking for Drug Discovery. Expert Opin. Drug Discov. 2015, 10 (11), 1189–1200.

(34) Koukos, P. I.; Xue, L. C.; Bonvin, A. M. J. J. Protein-Ligand Pose and Affinity Prediction: Lessons from D3R Grand Challenge 3. J. Comput. Aided Mol. Des. 2018.

(35) Hopkins, A. L.; Groom, C. R. The Druggable Genome. Nat. Rev. Drug Discov. 2002, 1, 727.

(36) Borhani, D. W.; Shaw, D. E. The Future of Molecular Dynamics Simulations in Drug Discovery. J. Comput. Aided Mol. Des. 2012, 26 (1), 15–26.

(37) Liu, X.; Shi, D.; Zhou, S.; Liu, H.; Liu, H.; Yao, X. Molecular Dynamics Simulations and Novel Drug Discovery. Expert Opin. Drug Discov. 2018, 13 (1), 23–37.

(38) Maximova, T.; Moffatt, R.; Ma, B.; Nussinov, R.; Shehu, A. Principles and Overview of Sampling Methods for Modeling Macromolecular Structure and Dynamics. PLoS Comput. Biol. 2016, 12 (4), e1004619.

(39) Mortier, J.; Rakers, C.; Bermudez, M.; Murgueitio, M. S.; Riniker, S.; Wolber, G. The Impact of Molecular Dynamics on Drug Design: Applications for the Characterization of Ligand-Macromolecule Complexes. Drug Discov. Today 2015, 20 (6), 686–702.

(40) Johnson, D. K.; Karanicolas, J. Druggable Protein Interaction Sites Are More Predisposed to Surface Pocket Formation than the Rest of the Protein Surface. PLOS Comput. Biol. 2013, 9 (3), e1002951.

(41) Harder, E.; Damm, W.; Maple, J.; Wu, C.; Reboul, M.; Xiang, J. Y.; Wang, L.; Lupyan, D.; Dahlgren, M. K.; Knight, J. L.; et al. OPLS3: A Force Field Providing Broad Coverage of Drug-like Small Molecules and Proteins. J. Chem. Theory Comput. 2016, 12 (1), 281–296.

(42) Totrov, M.; Abagyan, R. Flexible Ligand Docking to Multiple Receptor Conformations: A Practical Alternative. Curr. Opin. Struct. Biol. 2008, 18 (2), 178–184.

(43) Amaro, R. E.; Li, W. W. Emerging Methods for Ensemble-Based Virtual Screening. Curr. Top. Med. Chem. 2010, 10 (1), 3–13.

(44) Tarcsay, Á.; Paragi, G.; Vass, M.; Jójárt, B.; Bogár, F.; Keserű, G. M. The Impact of Molecular Dynamics Sampling on the Performance of Virtual Screening against GPCRs. J. Chem. Inf. Model. 2013, 53 (11), 2990–2999.

(45) Hazuda, D. J.; Anthony, N. J.; Gomez, R. P.; Jolly, S. M.; Wai, J. S.; Zhuang, L.; Fisher, T. E.; Embrey, M.; Guare, J. P.; Egbertson, M. S.; et al. A Naphthyridine Carboxamide Provides Evidence for Discordant Resistance between Mechanistically Identical Inhibitors of HIV-1 Integrase. Proc. Natl. Acad. Sci. U. S. A. 2004, 101 (31), 11233–11238.

(46) Pietrucci, F.; Vargiu, A. V.; Kranjc, A. HIV-1 Protease Dimerization Dynamics Reveals a Transient Druggable Binding Pocket at the Interface. Sci. Rep. 2015, 5, 18555.

(47) Schames, J. R.; Henchman, R. H.; Siegel, J. S.; Sotriffer, C. A.; Ni, H.; McCammon, J. A. Discovery of a Novel Binding Trench in HIV Integrase. J. Med. Chem. 2004, 47 (8), 1879–1881.

(48) Summa, V.; Petrocchi, A.; Bonelli, F.; Crescenzi, B.; Donghi, M.; Ferrara, M.; Fiore, F.; Gardelli, C.; Gonzalez Paz, O.; Hazuda, D. J.; et al. Discovery of Raltegravir, a Potent, Selective Orally Bioavailable HIV-Integrase Inhibitor for the Treatment of HIV-AIDS Infection. J. Med. Chem. 2008, 51 (18), 5843–5855.

(49) Zhong, H.; Carlson, H. A. Computational Studies and Peptidomimetic Design for the Human P53-MDM2 Complex. Proteins Struct. Funct. Bioinforma. 2005, 58 (1), 222–234.

(50) Li, Y. Y.; An, J.; Jones, S. J. A Computational Approach to Finding Novel Targets for Existing Drugs. PLoS Comput. Biol. 2011, 7 (9), e1002139.

(51) Lin, J.-H.; Perryman, A. L.; Schames, J. R.; McCammon, J. A. Computational Drug Design Accommodating Receptor Flexibility: The Relaxed Complex Scheme. J. Am. Chem. Soc. 2002, 124 (20), 5632–5633.

(52) Kuroda, D.; Gray, J. J. Pushing the Backbone in Protein-Protein Docking. Structure 2016, 24 (10), 1821–1829.

(53) Hamelberg, D.; McCammon, J. A. Accelerated Molecular Dynamics: A Promising and Efficient Simulation Method for Bio-molecules. J. Chem. Phys. 2004, 120 (24), 11919–11929.

(54) Luitz, M. P.; Zacharias, M. Protein-Ligand Docking Using Hamiltonian Replica Exchange Simulations with Soft Core Potentials. J. Chem. Inf. Model. 2014, 54 (6), 1669–1675.

(55) Sugita, Y.; Okamoto, Y. Replica-Exchange Molecular Dynamics Method for Protein Folding. Chem. Phys. Lett. 1999, 314 (1), 141–151.

(56) Laio, A.; Parrinello, M. Escaping Free-Energy Minima. Proc. Natl. Acad. Sci. 2002, 99 (20), 12562–12566.

(57) Grubmüller, H. Predicting Slow Structural Transitions in Macromolecular Systems: Conformational Flooding. Phys. Rev. E 1995, 52 (3), 2893–2906.

(58) Huber, T.; Torda, A. E.; van Gunsteren, W. F. Local Elevation: A Method for Improving the Searching Properties of Molecular Dynamics Simulation. J. Comput. Aided Mol. Des. 1994, 8 (6), 695–708.

(59) Antolin, A. A.; Carotti, A.; Nuti, R.; Hakkaya, A.; Camaioni, E.; Mestres, J.; Pellicciari, R.; Macchiarulo, A. Exploring the Effect of PARP-1 Flexibility in Docking Studies. J. Mol. Graph. Model. 2013, 45, 192–201.

(60) Bolia, A.; Ozkan, S. B. Adaptive BP-Dock: An Induced Fit Docking Approach for Full Receptor Flexibility. J. Chem. Inf. Model. 2016, 56 (4), 734–746.

(61) Miao, Y.; Goldfeld, D. A.; Moo, E. V.; Sexton, P. M.; Christopoulos, A.; McCammon, J. A.; Valant, C. Accelerated Structure-Based Design of Chemically Diverse Allosteric Modulators of a Muscarinic G Protein-Coupled Receptor. Proc. Natl. Acad. Sci. 2016, 113 (38), E5675–E5684.

(62) Motta, S.; Bonati, L. Modeling Binding with Large Conformational Changes: Key Points in Ensemble-Docking Approaches. J. Chem. Inf. Model. 2017, 57 (7), 1563–1578.

(63) Oleinikovas, V.; Saladino, G.; Cossins, B. P.; Gervasio, F. L. Understanding Cryptic Pocket Formation in Protein Targets by Enhanced Sampling Simulations. J. Am. Chem. Soc. 2016, 138 (43), 14257–14263.

(64) Osguthorpe, D. J.; Sherman, W.; Hagler, A. T. Generation of Receptor Structural Ensembles for Virtual Screening Using Binding Site Shape Analysis and Clustering. Chem. Biol. Drug Des. 2012, 80 (2), 182–193.

(65) Seeliger, D.; De Groot, B. L. Conformational Transitions upon Ligand Binding: Holo-Structure Prediction from Apo Conformations. PLoS Comput. Biol. 2010, 6 (1), e1000634.

(66) Uehara, S.; Tanaka, S. Cosolvent-Based Molecular Dynamics for Ensemble Docking: Practical Method for Generating Druggable Protein Conformations. J. Chem. Inf. Model. 2017, 57 (4), 742–756.

(67) Barducci, A.; Bussi, G.; Parrinello, M. Well-Tempered Metadynamics: A Smoothly Converging and Tunable Free-Energy Method. Phys. Rev. Lett. 2008, 100 (2), 020603.

(68) Piana, S.; Laio, A. A Bias-Exchange Approach to Protein Folding. J. Phys. Chem. B 2007, 111 (17), 4553–4559.

(69) Moréra, S.; Larivière, L.; Kurzeck, J.; AschkeSonnenborn, U.; Freemont, P. S.; Janin, J.; Rüger, W. High Resolution Crystal Structures of T4 Phage β-Glucosyltransferase: Induced Fit and Effect of Substrate and Metal Binding11Edited by R. Huber. J. Mol. Biol. 2001, 311 (3), 569–577.

(70) de Groot, B. L.; van Aalten, D. M. F.; Scheek, R. M.; Amadei, A.; Vriend, G.; Berendsen, H. J. C. Prediction of Protein Conformational Freedom from Distance Constraints. Proteins Struct. Funct. Bioinforma. 1997, 29 (2), 240–251.

(71) Seeliger, D.; Haas, J.; de Groot, B. L. Geometry-Based Sampling of Conformational Transitions in Proteins. Structure 2007, 15 (11), 1482–1492.

(72) Mlsna, D.; Monzingo, A. F.; Katzin, B. J.; Ernst, S.; Robertus, J. D. Structure of Recombinant Ricin A Chain at 2.3 Å. Protein Sci. 1993, 2 (3), 429–435.

(73) Yan, X.; Hollis, T.; Svinth, M.; Day, P.; Monzingo, A. F.; Milne, G. W. A.; Robertus, J. D. Structure-Based Identification of a Ricin Inhibitor. J. Mol. Biol. 1997, 266 (5), 1043–1049.

(74) Hartshorn, M. J.; Verdonk, M. L.; Chessari, G.; Brewerton, S. C.; Mooij, W. T. M.; Mortenson, P. N.; Murray, C. W. Diverse, High-Quality Test Set for the Validation of Protein−Ligand Docking Performance. J. Med. Chem. 2007, 50 (4), 726–741.

(75) Ravindranath, P. A.; Forli, S.; Goodsell, D. S.; Olson, A. J.; Sanner, M. F. AutoDockFR: Advances in Protein-Ligand Docking with Explicitly Specified Binding Site Flexibility. PLoS Comput. Biol. 2015, 11 (12), e1004586.

(76) Trott, O.; Olson, A. J. AutoDock Vina: Improving the Speed and Accuracy of Docking with a New Scoring Function, Efficient Optimization, and Multithreading. J. Comput. Chem. 2010, 31 (2), 455–461.

(77) Dominguez, C.; Boelens, R.; Bonvin, A. M. J. J. HADDOCK: A Protein−Protein Docking Approach Based on Biochemical or Biophysical Information. J. Am. Chem. Soc. 2003, 125 (7), 1731–1737.

(78) Morris, G. M.; Huey, R.; Lindstrom, W.; Sanner, M. F.; Belew, R. K.; Goodsell, D. S.; Olson, A. J. AutoDock4 and AutoDockTools4: Automated Docking with Selective Receptor Flexibility. J. Comput. Chem. 2009, 30 (16), 2785–2791.

(79) Meiler, J.; Baker, D. ROSETTALIGAND: Protein-Small Molecule Docking with Full Side-Chain Flexibility. Proteins Struct. Funct. Bioinforma. 2006, 65 (3), 538–548.

(80) Bakan, A.; Nevins, N.; Lakdawala, A. S.; Bahar, I. Druggability Assessment of Allosteric Proteins by Dynamics Simulations in the Presence of Probe Molecules. J. Chem. Theory Comput. 2012, 8 (7), 2435–2447.

(81) Leis, S.; Zacharias, M. Efficient Inclusion of Receptor Flexibility in Grid-Based Protein-Ligand Docking. J. Comput. Chem. 2011, 32 (16), 3433–3439.

(82) Pandini, A.; Fornili, A. Using Local States To Drive the Sampling of Global Conformations in Proteins. J. Chem. Theory Comput. 2016, 12 (3), 1368–1379.

(83) Pietrucci, F.; Laio, A. A Collective Variable for the Efficient Exploration of Protein Beta-Sheet Structures: Application to SH3 and GB1. J. Chem. Theory Comput. 2009, 5 (9), 2197–2201.

(84) Case, D.; Betz, R.; Cerutti, D.; Cheatham III, T.; Darden, T.; Duke, R. A. G., TJ; Gohlke, H.; Goetz, A.; Homeyer, N.; Izadi, S.; et al. Amber 16; 2016.

(85) Maier, J. A.; Martinez, C.; Kasavajhala, K.; Wickstrom, L.; Hauser, K. E.; Simmerling, C. Ff14SB: Improving the Accuracy of Protein Side Chain and Backbone Parameters from Ff99SB. J. Chem. Theory Comput. 2015, 11 (8), 3696–3713.

(86) Wang, J.; Wolf, R. M.; Caldwell, J. W.; Kollman, P. A.; Case, D. A. Development and Testing of a General Amber Force Field. J. Comput. Chem. 2004, 25 (9), 1157–1174.

(87) Frisch, M. J.; Trucks, G. W.; Schlegel, H. B.; Scuseria, G. E.; Robb, M. A.; Cheeseman, J. R.; Scalmani, G.; Barone, V.; Mennucci, B.; Petersson, G. A.; et al. Gaussian∼09 Revision E.01.

(88) Hopkins, C. W.; Le Grand, S.; Walker, R. C.; Roitberg, A. E. Long-Time-Step Molecular Dynamics through Hydrogen Mass Repartitioning. J. Chem. Theory Comput. 2015, 11 (4), 1864–1874.

(89) Abraham, M. J.; Murtola, T.; Schulz, R.; Páll, S.; Smith, J. C.; Hess, B.; Lindahl, E. GROMACS: High Performance Molecular Simulations through Multi-Level Parallelism from Laptops to Super-computers. SoftwareX 2015, 1-2, 19–25.

(90) Tribello, G. A.; Bonomi, M.; Branduardi, D.; Camilloni, C.; Bussi, G. PLUMED 2: New Feathers for an Old Bird. Comput. Phys. Commun. 2014, 185 (2), 604–613.

(91) Sousa da Silva, A. W.; Vranken, W. F. ACPYPE - Ante-Chamber PYthon Parser InterfacE. BMC Res. Notes 2012, 5 (1), 367.

(92) Laio, A.; Rodriguez-Fortea, A.; Gervasio, F. L.; Ceccarelli, M.; Parrinello, M. Assessing the Accuracy of Metadynamics. J. Phys. Chem. B 2005, 109 (14), 6714–6721.

(93) Celebi, M. E.; Kingravi, H. A.; Vela, P. A. A Comparative Study of Efficient Initialization Methods for the K-Means Clustering Algorithm. Expert Syst. Appl. 2013, 40 (1), 200–210.

(94) van Zundert, G. C. P.; Rodrigues, J. P. G. L. M.; Trellet, M.; Schmitz, C.; Kastritis, P. L.; Karaca, E.; Melquiond, A. S. J.; van Dijk, M.; de Vries, S. J.; Bonvin, A. M. J. J. The HADDOCK2.2 Web Server: User-Friendly Integrative Modeling of Biomolecular Complexes. Comput. Resour. Mol. Biol. 2016, 428 (4), 720–725.

(95) Kurkcuoglu, Z.; Koukos, P. I.; Citro, N.; Trellet, M. E.; Rodrigues, J. P. G. L. M.; Moreira, I. S.; Roel-Touris, J.; Melquiond, A. S. J.; Geng, C.; Schaarschmidt, J.; et al. Performance of HADDOCK and a Simple Contact-Based Protein-Ligand Binding Affinity Predictor in the D3R Grand Challenge 2. J. Comput. Aided Mol. Des. 2018, 32 (1), 175–185.

(96) Schrödinger Release 2015-4: Maestro; Schrödinger, LLC: New York, NY, 2015.

(97) Humphrey, W.; Dalke, A.; Schulten, K. VMD: Visual Molecular Dynamics. J. Mol. Graph. 1996, 14 (1), 33–38.

